# Cohesin and condensin extrude loops in a cell-cycle dependent manner

**DOI:** 10.1101/821306

**Authors:** Stefan Golfier, Thomas Quail, Hiroshi Kimura, Jan Brugués

## Abstract

Chromatin undergoes a dramatic reorganization during the cell cycle^1–3^. In interphase, chromatin is organized into compartments and topological-associating domains (TADs) that are cell-type specific^4–7^, whereas in metaphase, chromosomes undergo large-scale compaction, leading to the loss of specific boundaries and the shutdown of transcription^8–12^. Loop extrusion by structural maintenance of chromosomes complexes (SMCs) has been proposed as a mechanism to organize chromatin in interphase and metaphase^13–19^. However, the requirements for chromatin organization in these cell phases are very different, and it is unknown whether loop extrusion dynamics and the complexes that extrude them also differ. Here, we used *Xenopus* egg extracts to reconstitute and image loop extrusion of single DNA molecules during the cell cycle. We show that loops form in both metaphase and interphase, but with distinct dynamic properties. Condensin extrudes asymmetric loops in metaphase, whereas cohesin extrudes symmetric loops in interphase. Our data show that loop extrusion is a general mechanism for the organization of DNA, with dynamic and structural properties that are molecularly regulated during the cell cycle.

## Main text

To visualize loop formation in *Xenopus laevis* egg extracts, we attached lambda-phage DNA to a cover slip using biotin-streptavidin linkers^20^ in a custom-built microfluidic chamber (Fig. 1A). Addition of either metaphase-arrested or interphase *Xenopus* egg extracts into the chamber triggered the formation of small DNA enrichments, consistent with nucleosomal deposition^21,22^, that rapidly reduced any slack in the DNA molecules (Fig. S1A and Movies 1-2). To allow enough slack for the formation of loops, we abolished nucleosomal assembly along the strand by depleting ~90-95% of soluble H3-H4 heterodimers in the extract^23^ (Fig. S1B). This led to the formation of compacted DNA clusters that grew in size over time both in metaphase and interphase extracts (Fig. 1B and Movies 3-8). To investigate whether these clusters exhibit a topology consistent with DNA loops, we applied hydrodynamic forces to the DNA strand by introducing a flow in the perpendicular direction to the strand. This procedure revealed DNA clusters with a characteristic loop topology similar to loop extrusion by yeast condensin *in vitro*^24^ (Fig. 1C, Fig. S1C and Movie 9-12). In mock-depleted extracts, loops also formed but at a much lower frequency (Fig. S1D and Movie 13) and seemed to compete with nucleosomes for free DNA. These results show that single DNA loop extrusion can be reconstituted in *Xenopus* egg extracts in metaphase and interphase.

**Figure 1:**
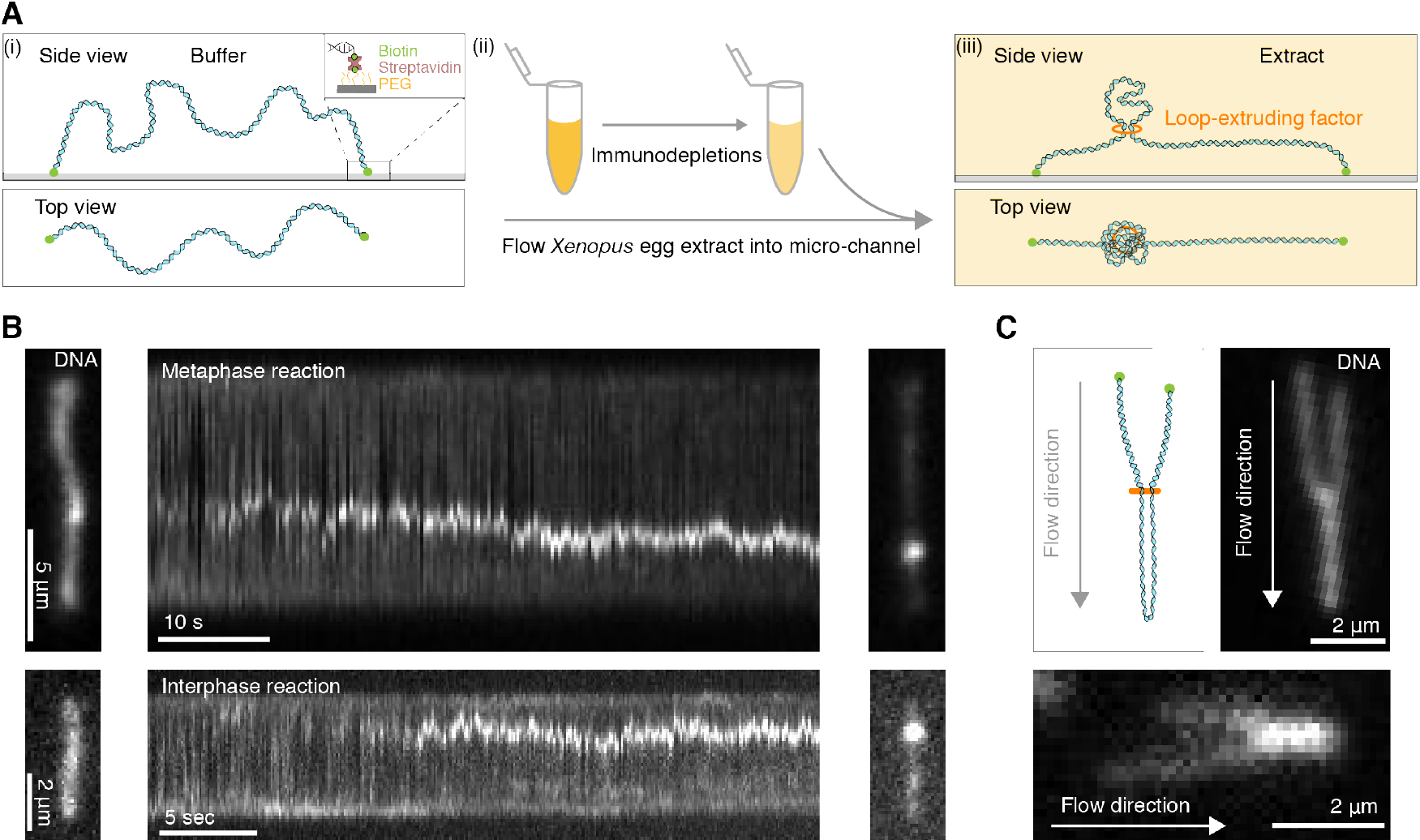
Single DNA molecule assay for direct visualization of DNA looping in *Xenopus* egg extracts. (A) (i) Side and top view schematics of a single strand of λ-phage DNA attached to a functionalized cover slip via biotin-streptavidin linkers. (ii) *Xenopus* egg extract is flowed into the microfluidic chamber. (iii) Side and top view schematics visualizing how soluble active loop-extruding factors extrude loops in nucleosome-depleted extract. (B) Dynamics of the formation of DNA loops induced by nucleosome-depleted extract in metaphase (upper) and interphase (lower). Snapshot of a single molecule of λ-phage DNA visualized using Sytox Orange *in vitro* preceding treatment with nucleosome-depleted extract (left). Kymograph of DNA signal over time displaying a looping event upon addition of nucleosome-depleted extract (middle). Snapshot of steady-state DNA looping event after ~40 sec (right). (C) Hydrodynamic flows reveal loop topology within DNA cluster. (i) Schematic of the loop topology revealed upon flow. (ii) Topology of extract-induced DNA loops in metaphase (upper) and interphase (lower) visualized using Sytox Orange revealed upon flow in the direction of the arrow.

Current theoretical models propose that SMCs organize chromatin by extruding symmetric DNA loops^14,16,18^. However, recent experimental studies have shown that yeast condensin can extrude loops asymmetrically *in vitro*^24^. Although this supports the loop-extrusion hypothesis, it is inconsistent with the symmetric extrusion predicted by theory^25,26^. One reason for this discrepancy could be that the properties of loop extrusion in cellular contexts differ from those *in vitro*. To characterize the dynamic properties of loop formation in *Xenopus* extracts, we quantified the DNA distribution inside the loop and to the left and right of the loop as a function of time (Fig. 2). This allows the rate of loop extrusion to be obtained from the amount of DNA that goes into the loop over time, and to determine whether this amount of DNA is extruded from one or both of the outer non-looped regions. Briefly, we summed the fluorescence intensity of the DNA along the perpendicular direction to the stretched DNA strand and tracked the loop position defined by the local maximum of this DNA intensity. We then fitted a Gaussian function to the loop region and defined the loop boundaries as ±2 standard deviations away from the maximum value of the fit (Fig. 2A). We obtained the amount of DNA inside the loop as the difference between the integrated intensity in the loop region minus the offset intensity from the Gaussian fit.

**Figure 2:**
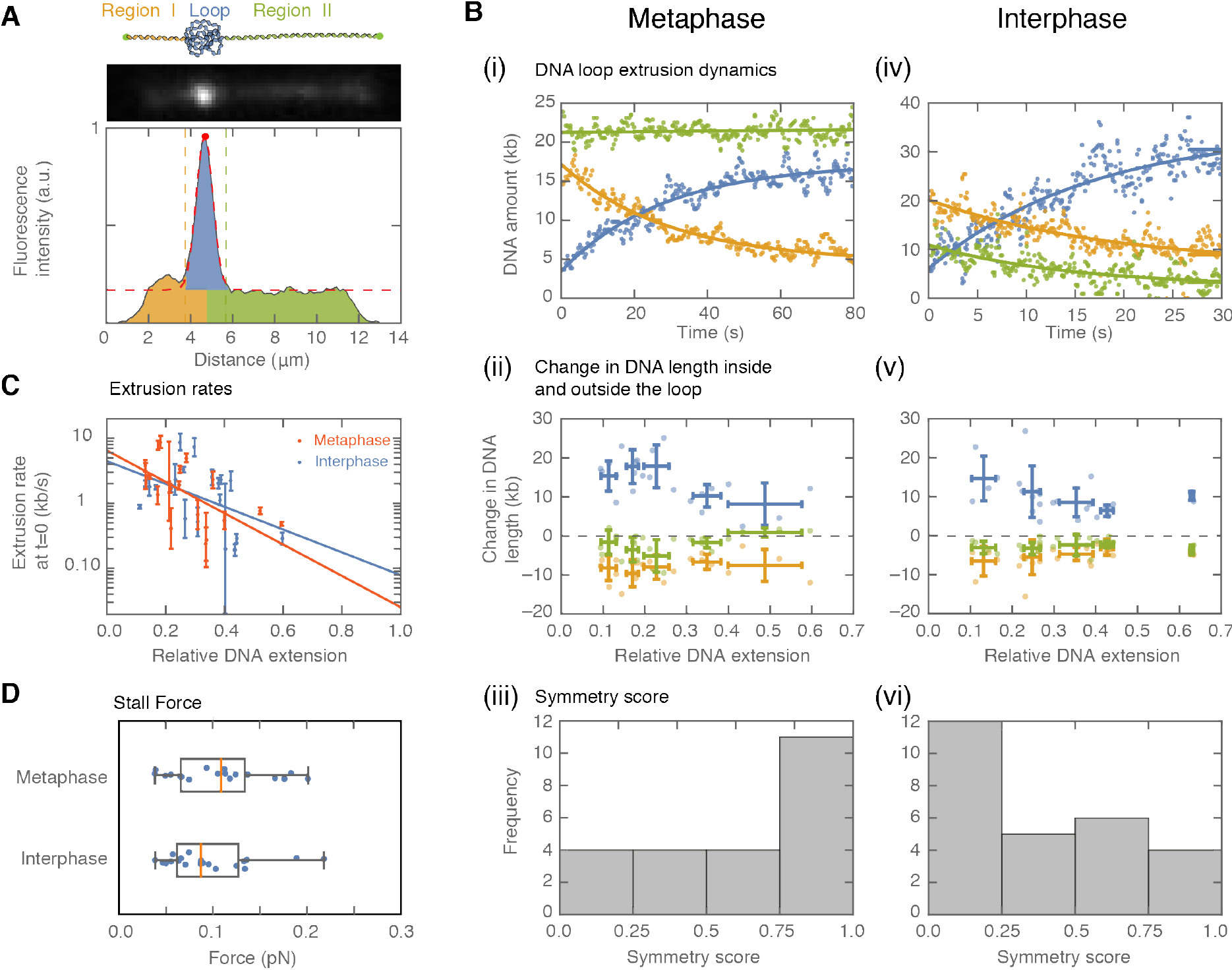
Symmetry of loop extrusion is cell-cycle dependent with similar extrusion rates and stalling forces. (A) *Upper*: Schematic of the top view of a DNA-looping event segmented into three regions: region I (orange), region II (green), and the loop region (blue). *Middle*: Snapshot of DNA-looping event where DNA is labelled using Sytox Orange. *Bottom*: The integrated fluorescence intensity of the DNA generated by summing the intensity values along the perpendicular axis of the strand. The dashed red line represents a Gaussian fit to the data. Signal values above the fit’s offset define the looped region given in blue; signal values below this threshold correspond to the non-looped regions I and II, given in orange in green. (B) Dynamics of DNA looping in metaphase and interphase. (Bi,iv) DNA amount as a function of time computed for the looped region (blue), and regions I and II (green and orange). The dots represent experimental data and the solid lines represent exponential fits to the data. (Bii, v) Total change in the amount of DNA for the looped region and non-looped regions I and II plotted as a function of relative DNA extension. Error bars correspond to standard deviations of data clustered by proximity. Points represent raw data. (Biii, vi) Analysis of loop extrusion symmetry shows asymmetric extrusion (symmetry score ~ 1) in metaphase (Biii) and symmetric extrusion (symmetry score ~ 0) in interphase (Bvi). (C) Growth rates of DNA loop extrusion in metaphase (orange) and interphase (blue) as a function of relative DNA extension. Error bars are obtained from error propagation of the uncertainties of the exponential fit parameters. (D) Box plots of the stall forces for DNA loop extrusion in metaphase and interphase.

Finally, the amount of DNA to the left and right of the loop corresponds to the integrated intensity of the DNA strands outside the loop region (see Methods). This assay allows us to observe loop extrusion in extract, to quantify the partitioning of DNA between the looped and outer regions, and to examine the symmetry of the reeling-in of DNA. When applied to nucleosome-depleted extract arrested in metaphase, this assay showed that loops in metaphase are extruded at 2.3 ± 0.5 kb/s (mean ± SEM) and appeared to be extruded from one side of the loop (Fig. 2Bi, 2C, and Fig. S2).

To further characterize the symmetry of extrusion in metaphase, we quantified the total decrease in DNA from the left and right regions of the loop between the onset of loop formation and the final steady-state size of the loop (Fig. 2Bii). We used these quantities to define a symmetry score as the relative difference between the decrease of these two regions and the total amount of extruded DNA. The histogram of symmetry scores peaked at 1, which corresponds to asymmetric loops (Fig. 2Biii). Our assay allows the final steady state amount of DNA in the loop to be determined as a function of the end-to-end DNA-binding distance. These measurements show that the loop-extrusion process stops when the relative extension of DNA outside of the loop reaches ~75%, with a corresponding stall force of 0.41 pN ± 0.01 pN (mean ± SEM), (Fig. 2D). Thus, our analysis demonstrates that loop extrusion in metaphase is preferentially one-sided, with extrusion speeds and stall forces similar to those measured *in vitro*^24,27,28^.

Next, we used nucleosome-depleted extract in interphase to investigate whether the dynamics of loop extrusion share similar properties during the cell cycle (Fig. 2Civ-vi). Loop extrusion in interphase displayed a similar distribution of extrusion rates of 2.1 ± 0.5 kb/s, and stall forces, 0.23 pN ± 0.01 pN, (Fig. 2C, 2D). However, the distribution of symmetry scores of these loops peaked towards zero, suggesting that these loops are preferentially extruded symmetrically. Thus, the mechanisms of loop extrusion differ between interphase and metaphase.

The different dynamic properties of loop formation we observe in interphase and metaphase suggest that different molecular activities may be responsible for loop formation during the cell cycle^15,29^. Recent work has suggested that cohesin could extrude loops in interphase, though this activity has not been directly visualized in cellular contexts^30–32^. Thus, cell-cycle-dependent activities of condensin and cohesin could account for the transition between symmetric and non-symmetric loop extrusion^15^. To assess the role of cohesin and condensin during loop extrusion in interphase and metaphase, we selectively depleted these proteins in egg extract using antibodies (Fig. S3). We then tested for loop extrusion activity in each depleted condition. We found that, in metaphase, the occurrence of loop extrusion was significantly (p<0.01) reduced upon depletion of condensin I and II, but was unaffected by cohesin depletion (Fig. 3A). In contrast, there was a significant (p<0.01) decrease in loop extrusion following cohesin depletion in interphase, but was unaffected by condensin depletion (Fig. 3A). We confirmed these depletions with immunostainings that showed colocalization of cohesin and condensin with the loops observed in interphase and metaphase, respectively (Fig. 3B). Thus, cohesin extrudes symmetric loops during interphase, whereas condensin extrudes asymmetric loops in metaphase.

**Figure 3:**
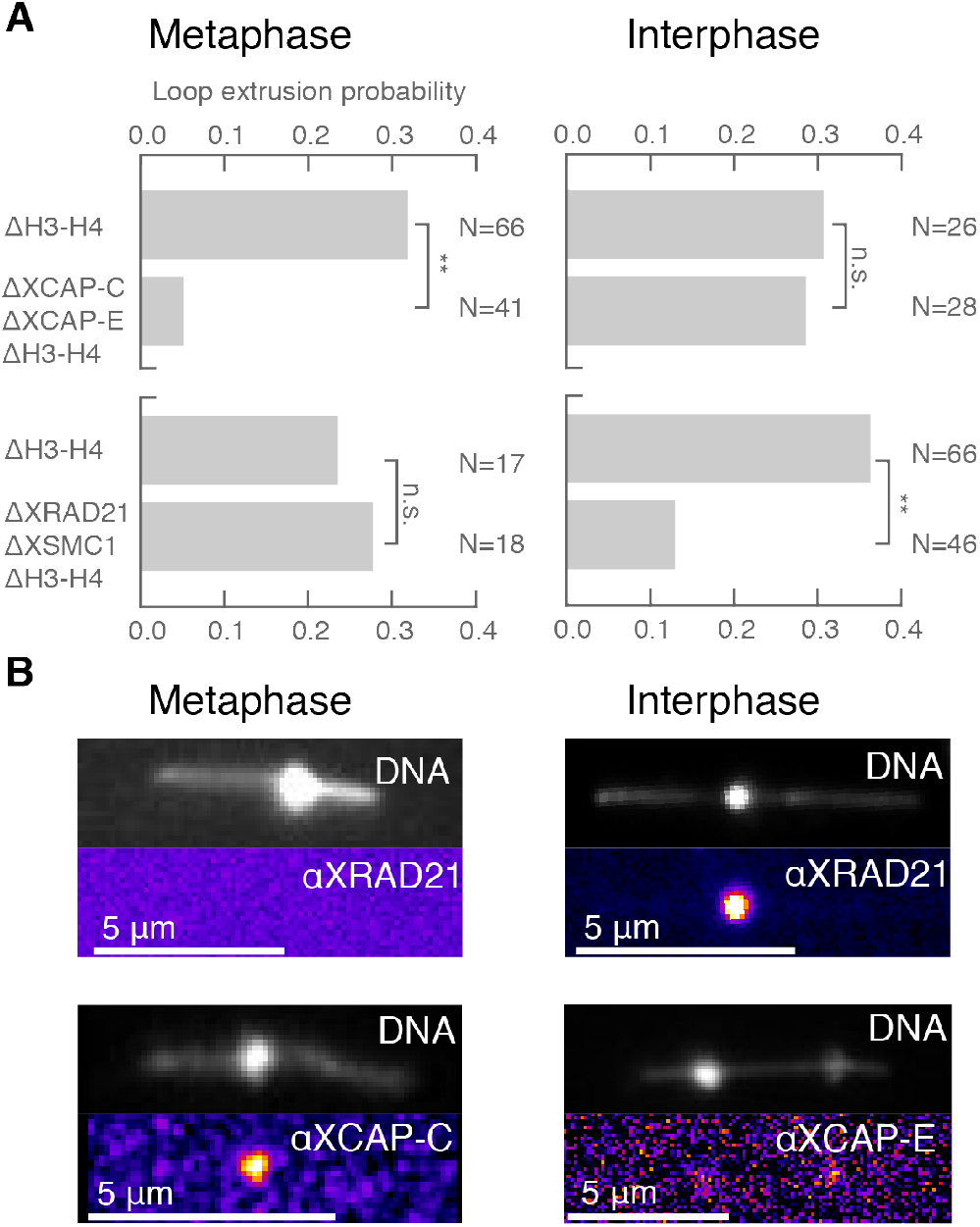
Condensin extrudes loops in metaphase and cohesin extrudes loops in interphase. (A) Loop extrusion probability in metaphase and interphase under different depletion conditions. In metaphase, co-depleting condensin I, condensin II, and H3-H4 (using anti-XCAP-C/E and anti-H4K12Ac) significantly (p<0.01, Binomial test) reduced loop extrusion probability compared to H3-H4-depleted extract, but had no effect in interphase. In contrast, depleting cohesin and H3-H4 (using anti-XRAD21/XSMC1 and anti-H4K12Ac) significantly (p<0.01) decreased loop extrusion probability in interphase but had no effect in metaphase. (B) Snapshots of antibody stainings of representative loops in metaphase and interphase.

Our findings provide the first evidence that loop extrusion is a general mechanism of DNA organization in a cellular context, and furthermore, that it is differentially regulated during the cell cycle. This regulation is achieved by the distinct activities of cohesin^30,31^ and condensin^33,34^ during interphase and metaphase, and may control different levels of DNA organization during the cell cycle: from chromatin that is mostly decondensed and spatially organized into TAD structures during interphase to highly compacted chromosomes in metaphase. Symmetric loop extrusion by cohesin in interphase may ensure the formation of specific TAD boundaries by bringing together neighboring CTCF sites^35–37^. In metaphase, reorganization of chromosomes into compact chromatids requires the loss of boundaries and shut down of transcription^3,8,12^, which may be achieved by condensin spooling activity. However, our findings highlight the need to revise our understanding of how loop extrusion accounts for the different levels of chromatin organization in interphase and metaphase, and why different extrusion symmetries are required during the cell cycle. Our assay will allow dissection of the molecular components that regulate the dynamic properties of loop formation in physiological contexts, and make it possible to reconstitute more complex processes such as the formation of boundary elements and the interplay between transcription, replication, and loop extrusion.

## Methods

### *Xenopus laevis* egg extract preparation and immunodepletion

Cytostatic factor (CSF)-arrested *Xenopus laevis* egg extract was prepared as described previously^38^. In brief, unfertilized oocytes were dejellied and crushed by centrifugation, generating an extract that was arrested in meiosis II. We added protease inhibitors (LPC: leupeptin, pepstatin, chymostatin) and cytochalasin D (CyD) to a final concentration of 10 μg/ml each to the extract. In order to generate interphase extracts, CaCl_2_ was added to a final concentration of 0.4 mM. To immunodeplete soluble H3-H4 heterodimers from the extract^23,39^, we coupled 130 μg of a mouse monoclonal anti-H4K12Ac (gift from Hiroshi Kimura) to 12.5 μl rProtein A Sepharose (GE Healthcare) slurry in antibody coupling buffer (10 mM K-HEPES pH=8, 150 mM NaCl), rotating overnight at 4 °C. After several washes with a wash buffer (10 mM HEPES pH=7.7, 100 mM KCl, 150 mM Sucrose, 1 mM MgCl_2_), we combined 50 μl fresh CSF extract with the beads and incubated the bead-extract mixture for 1.5 hours on ice, occasionally flicking the tubes in order to prevent the beads settling to the bottom. After recovering the extract from the beads, we immediately proceeded with the experiment. We generated mock-depleted extracts with the same protocol using 130 μg random mouse IgG antibodies in 50 μl of fresh CSF extract. To co-deplete H3-H4 and both condensin I and condensin II, we coupled 130 μg anti-H4K12Ac and 10 μg rabbit polyclonal antibodies of both anti-XCAP-C and anti-XCAP-E to 15 μl rProtein A Sepharose slurry and performed the same H3-H4 depletion method. To co-deplete H3-H4 and cohesin, we coupled 130 *μg* anti-H4K12Ac and 10 μg rabbit polyclonal anti-XRad21 and 10 μg anti-XSMC1 to 15 μl rProtein A Sepharose and performed the same H3-H4 depletion method.

### Western blots

We prepared 1:25 dilutions of immunodepleted extract in 1X sample loading buffer (50 mM Tris-HCl, pH=6.8, 2% SDS, 10% glycerol, 0.006% bromophenol blue, 100 mM DTT), ran gel electrophoresis on a gradient gel, transferred to a nitrocellulose membrane with a semi-dry transfer approach, and performed primary antibody incubation with polyclonal rabbit antibodies anti-H3 (1:10000, ab1791), anti-XSMC1 (1:2500, MPI-CBG antibody facility), anti-XCAP-C (1:2000, MPI-CBG antibody facility) and monoclonal mouse antibodies to detect tubulin using anti-DM1a (1:10000, MPI-CBG antibody facility). We detected primary antibodies using LI-COR IRDye secondary antibodies and imaged the western blots on an Odyssey Infrared Imaging System. We analyzed the blots using FIJI.

### Antibody production and labeling

We raised rabbit polyclonal antibodies for immunodepletion against peptides SDIVATPGPRFHTV and DLTKYPDANPNPND corresponding to antibodies that target cohesin’s XRAD21 and XSMC1 subunits. We also raised rabbit polyclonal antibodies against peptides AAKGLAEMQSVG and SKTKERRNRMEVDK corresponding to antibodies that target XCAP-C and XCAP-E for both condensin I and II for immunodepletion^34^. We added a cysteine residue on the peptide’s N-terminus for sulfhydryl coupling, and subsequent keyhole limpet hemocyanin conjugation and affinity purification was performed by MPI-CBG antibody facility. We labeled antibodies with fluorophores for localization following the small-scale on-resin labeling technique from^40^. Briefly, we prepared a 200-μl pipette tip to act as our resin bed. We then loaded 40 μl of rProtein A Sepharose (GE Healthcare) resin into the tip, washing three times with 10 mM K-HEPES (pH=7.7), 150 mM NaCl. We labeled both the antibody targeting the cohesin subunit XRad21 and the antibody targeting condensin I and II’s subunit XCAP-C. We flowed 70 μg antibody 5 times consecutively through the packed resin bed in order to bind the antibody to the resin. The resin was then washed three times with 200 mM K-HEPES (pH=7.7). We then added 0.5 μl 50 mM NHS-Ester-Alexa488 (Alexa Fluor™ NHS Ester, A20000, Thermo Fischer) to 25 μl 200 mM K-HEPES (pH=7.7), and immediately added it to the resin, incubating the resin, antibody, and dye for 10-60 minutes at room temperature. To remove the unbound dye, the resin bed was washed 5 times with 10 mM K-HEPES (pH=7.7), 150 mM NaCl. We eluted the labelled antibody with 5×15 μl of 200 mM acetic acid. We neutralized each eluate immediately with 5 μl of 1 M Tris-HCl, pH=9, and cooled to 0 °C. The labelled antibody is stable for months kept at 4 °C.

### DNA functionalization

To biotinylate DNA purified from λ-phage (λ-DNA)^41^, we combined 10 μg of λ-DNA (NEB, N3011S) and 5 μl of a 10X polymerase buffer (50 mM Tris-HCl, pH=7.2, 10 mM MgSO4, 100 μM DTT) to a total reaction volume of 50 μl. We then heated the mixture up to 65 °C for 7 minutes to break apart the λ-DNA’s sticky ends. After heat treatment, we added 100x molar excess of biotinylated dATP, biotinylated dUTP, and dGTP, and dCTP. We then added 1 unit (~1 μl) of Klenow enzyme, mixed well, and incubated overnight at room temperature. We purified the biotinylated λ-DNA using ethanol precipitation and stored at −20 °C.

### PEGylation of cover slips and DNA micro-channel preparation

We functionalized glass cover slips with mPEG and PEG-Biotin. Briefly, we sonicated coverslips first in acetone for 15 minutes followed by 5 rinses with MilliQ water, and then another sonication step in 5 M KOH for 40 minutes. After rinsing the coverslips 3 times with water and then 3 times with methanol, we dried the coverslips with N_2_. We silanized the coverslips combining 250 ml methanol, 12.5 ml acetic acid, and 2.5 ml (3-aminopropyl)-trimethoxysilane, incubating the coverslips in this mixture for 10 minutes at room temperature, sonicating for 1 minute, and then incubating the coverslips for an additional 10 minutes. Next we rinsed the coverslips once with methanol, once with water, and once again methanol, and dried with N_2_. Then we mixed 100 mg mPEG and ~1.5 mg Biotin-PEG with 450 μl PEGylation buffer (0.1M Sodium Bicarbonate, pH=8.5), and spun the reaction at 10000 RPM for 1 minute. We pipetted 25 μl of the PEG mixture onto a dried, silanized coverslip and put another coverslip on top, generating a coverslip sandwich. We incubated these sandwiches over night in distilled water-filled pipette tip-boxes in the dark. After incubation, we carefully disassembled the coverslips, rinsed with MilliQ water, and dried with N_2_. To generate a channel for imaging, we first drilled holes through a cleaned cover slide—these holes acted as channel inlets and outlets. We placed custom-designed, laser-cut double-sided tape onto the coverslip, defining the channel geometry. We then placed a functionalized PEG-biotinylated coverslip on top of the double-sided tape, sealing the channel on either end with Valap. We filled the channel with ~10-15 μl of 0.1 mg/ml free streptavidin, incubating the channel with streptavidin for 1 minute. To remove the free, unbound streptavidin, we flushed ~100 μl channel washing buffer (40 mM Tris-HCl, pH=8.0, 20 mM NaCl, 0.4 mM EDTA) through the channel, using the drilled holes as channel inlets and outlets. We added 20 μl of 1:1000 biotinylated λ-DNA (~5 pM), incubating it for ~1 min and then washed the channel with 3×100 μl of channel washing buffer.

### Imaging

For live imaging of looping events, we fluorescently stained immobilized DNA strands with 50nM Sytox Orange (S11368, ThermoFisher), a DNA intercalating dye, in imaging buffer (50mM Tris-HCl pH 7.7, 50mM KCl, 2.5mM MgCl_2_, 2mM ATP) similar to^24^ or Xenopus Buffer (XB: 100mM KCl, 1mM MgCl_2_, 0.1mM CaCl2, 2mM ATP). We excited Sytox Orange-labelled DNA using a 561nm laser, and imaged the strands using a Nikon Eclipse microscope stand with a Nikon 100x/NA 1.49 oil SR Apo TIRF and an Andor iXon3 EMCCD camera using a frame-rate of 100 – 300ms. A highly inclined and laminated optical sheet (HILO) microscopy mode was established using a Nikon Ti-TIRF-E unit mounted onto the microscope stand to improve signal-to-noise ratio by cutting off background fluorescence signal from unbound DNA dye in the buffer. To trigger the formation of DNA loops, we flowed about 2ul of nucleosome depleted extract into the channel (total channel volume ~10ul) and let the extract diffuse further down the channel. We then imaged looping events at the moving front of the diffusing extract.

### Hydrodynamic stretching of loops

To visualize DNA loop topology which cannot be observed in the normal mode of data acquisition, we hydrodynamically stretched DNA strands that exhibited looping events using a flow-controlled syringe pump (Pro Sense B.V., NE-501), see also Movies 9-12. The flow direction was set to be perpendicular to the strand orientation by a cross-shaped channel design. Depending on the width of the channel, we used flow rates between 100 μl/min and 500 μl/min to extend DNA loops. Specifically, we introduced nucleosome depleted extract into the channel as described above and, upon loop formation, stretched DNA strands by flowing imaging buffer from the opposite side.

### Loop extrusion analysis

DNA traces were analyzed using custom-written Python scripts motivated by^24^. We converted movies of fluorescent DNA molecules into one-dimensional intensity profiles by summing the intensity values along the direction perpendicular to the DNA strand in each frame. We removed the background signal using a median filter. From the summed intensity profile for each frame we built kymographs by concatenating all time points (Fig. 2 and Fig. S1). To yield the amount of DNA inside and outside the loop for each time point, we segmented the DNA intensity profiles into a loop region and two regions outside of the loop by first finding the maximum intensity value as the position of the loop and subsequent fitting of a Gaussian around that position. We defined the boundaries of the loop region and the regions outside of the loop by the positions +/− 2x standard deviations from the center of the Gaussian fit. Summing the intensity values of the regions outside of the loop and integrating the intensity under the Gaussian fit yielded the proportions of total signal intensity in each of the three regions for each time point. The difference between the integrated intensity below the loop and the offset from the gaussian fit (corresponding to the intensity outside of the loop) was equally distributed to the regions outside of the loop as the signal from the incoming and outgoing DNA strands that are not part of the loop itself (Fig. 2A).

We calculated the relative sizes of the three regions in kilo-base pairs (kb) for each time frame by multiplying the 48.5 kb total length of lambda DNA with the ratio of each summed intensity value and the total summed intensity of the strand for every time point. From these values we calculated the relative change in each region over time by subtracting the averaged ten last data points from the averaged ten first data points in each region. We used the resulting values *a* and *b* for the region left and right of the loop to assign a symmetry score for each looping event by calculating

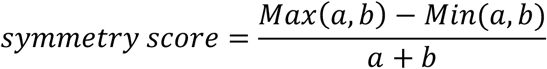

This procedure orders the extrusion from region *a* and *b* such that the symmetry score is always positive and ranges from 0 to 1. Our symmetry score intends to quantify the amount of DNA extruded into the loop from the outer regions. A positive relative change from one side implies that the no DNA from that side has been extruded into the loop, and thus we set that change to 0 (if a >0: a=0; if b >0: b=0).

We extracted the initial loop extrusion rates from the first derivative at time point zero of a single exponential fit to the values of the loop growth over time (Fig. 2B-C). The size of the loop at each time point further allowed us to calculate the extension of the DNA molecule by dividing the distance of the two immobilized ends of the DNA strand on the slide by the total length of the regions outside of the loop. From these values, we estimated the tension on the DNA strand for each time point by applying the Worm Like Chain Model of DNA^42^ to the extension of the DNA molecule. We determined the stall force of loop extrusion using the average of the last ten tension values per looping event when loop size reached steady state. For the analysis of extrusion stall forces we only used DNA strands where the loop extrusion did not end (or was stalled) at the DNA end-binding sites (N=25).

To quantify the effect of cohesin and condensin depletion, we determined the probability of loop extrusion by counting the number of observable loop extrusion events in all data taken for one condition and dividing it by the total number of DNA strands with sufficient slack (< 70% relative extension) to support the formation of a loop for that condition.

## Acknowledgements

We acknowledge A A Hyman, P. Tomancak, J. Guck, M. Loose, K. Ishihara, I. Patten, M. Sarov, M. Elsner and V. Murugesan for discussions and revision of the manuscript. We thank H. Andreas for frog maintenance, the Light Microscopy Facility (LMF), and the Antibody Facility at MPI-CBG.

## Contributions

S.G. designed the research, performed experiments and analyzed the data. T.Q. designed the research, performed experiments and wrote the manuscript. H.K. produced reagents. J.B. conceived and oversaw the project, designed the research, performed experiments, analyzed data and wrote the manuscript.

## Supplementary figures

**Supplementary Figure 1:**
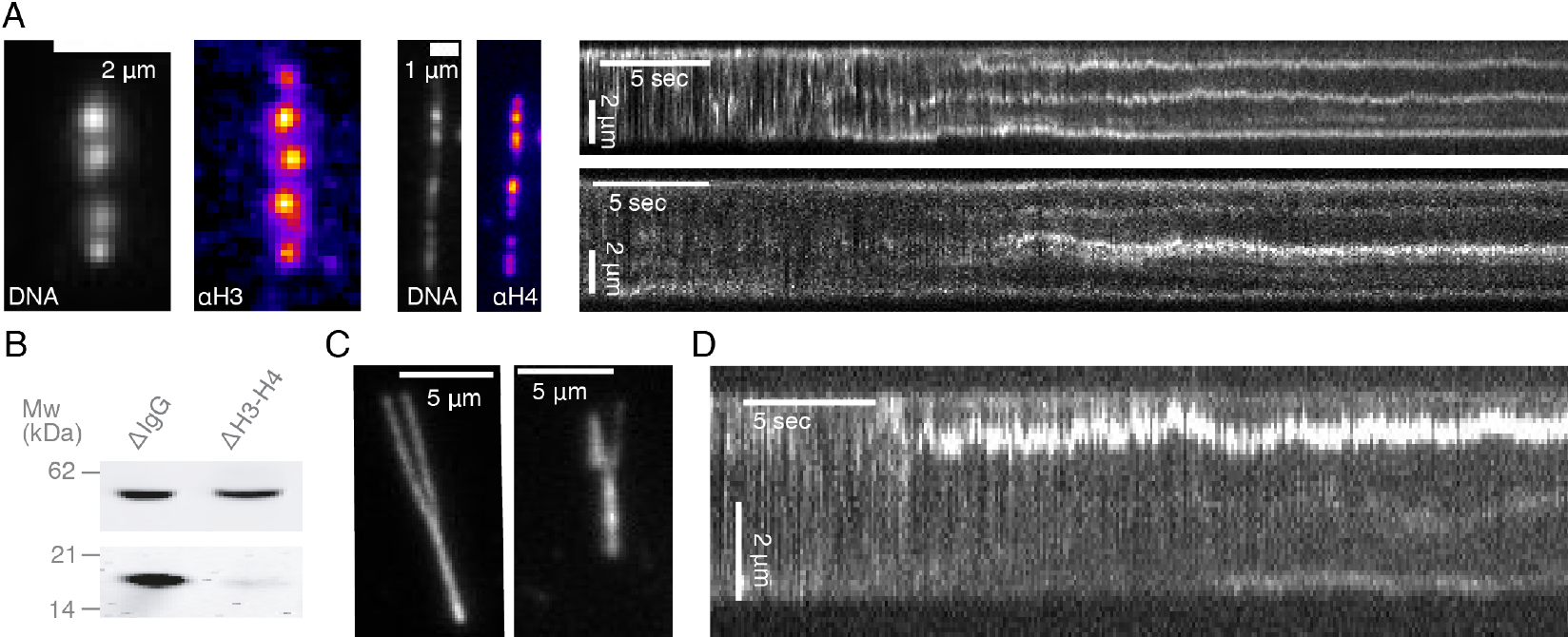
Characterization of cell-cycle-dependent looping dynamics in *Xenopus* egg extracts. (A) Addition of crude extract to strands of λ-phage DNA leads to the generation of multiple highly-enriched DNA clusters, suggestive of nucleosomal formation along the strand. Alexa488-labeled anti-H3 and anti-H4k12ac localize to these DNA clusters (left). Kymographs of nucleosomal cluster formation in both metaphase (upper) and interphase (lower) along a strand upon addition of crude extract. See also Movie 1 and 2. (B) Western blot showing approximately 90-95% depletion of soluble H3-H4 heterodimers. (C) Examples of two stretched loops in metaphase (left) and interphase (right) upon hydrodynamic pushing forces. (D) Kymograph of a looping event on a strand upon treatment with crude extract. See also Movie 13.

**Supplementary Figure 2:**
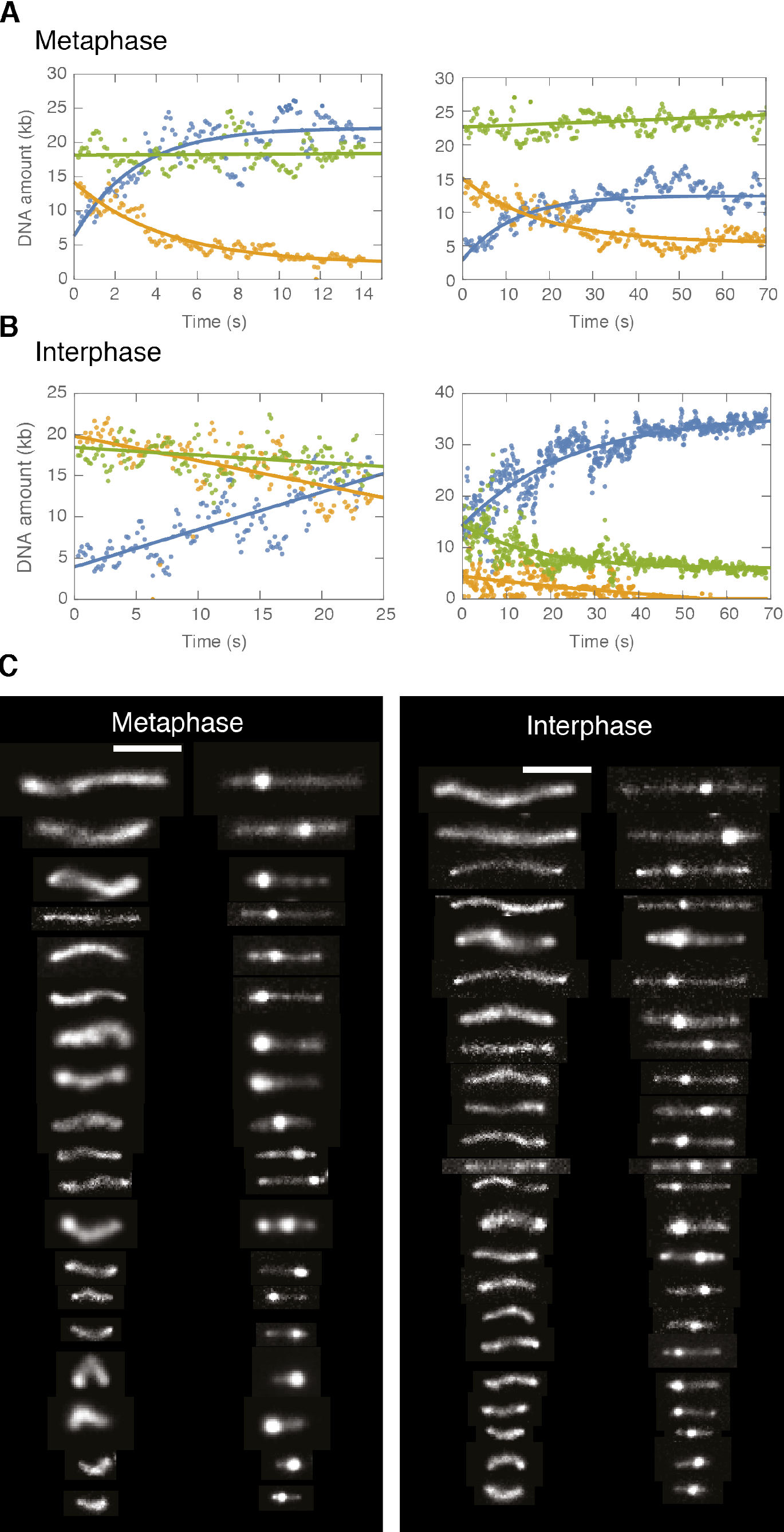
DNA looping examples demonstrating asymmetric looping in metaphase and symmetric looping in interphase. We tracked the position of a loop, and integrated the intensity of the loop as a function of time (given in blue). The green and orange colors correspond to the regions that are not in the loop. The dots are data from the analysis and the lines represent single exponential fits to the data. (A) In metaphase, we observed a saturating exponential increase in the DNA amount in the looped region, and only a single outer non-looped region decreasing in DNA amount, suggestive of asymmetric loop extrusion. (B) In interphase, we observed a saturating exponential increase in the DNA amount in the looped region, whereas both non-looped regions decreased in DNA amount, suggestive of symmetric loop extrusion. (C) Representative examples of DNA looping in nucleosome-depleted interphase and metaphase extract ordered by initial slack. Left image represents the initial DNA configuration and right image the corresponding final loop configuration. Scale bar is 5 μm.

**Supplementary Figure 3:**
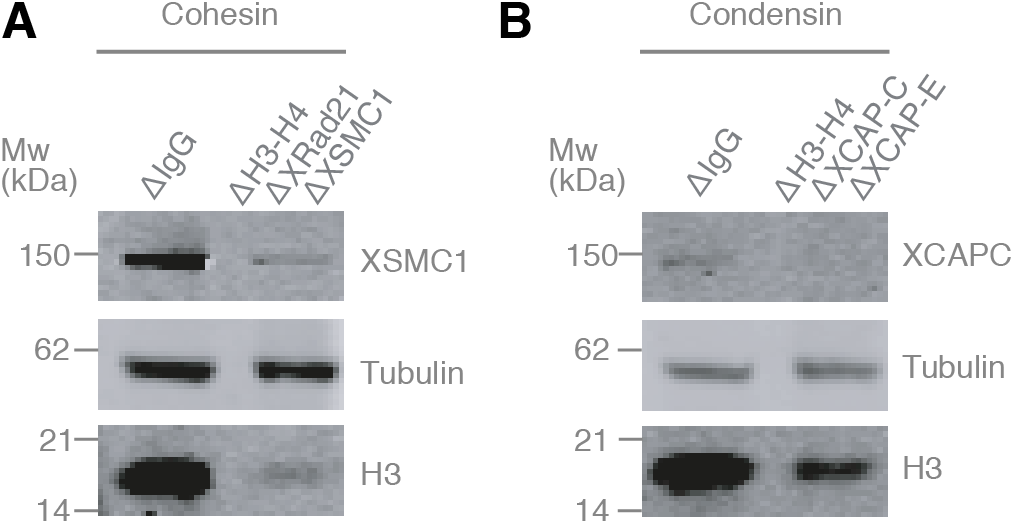
Co-immunodepletions of *Xenopus* egg extracts using antibodies targeting H3-H4, cohesin, and condensin I and II. (A) Co-immunodepletion of H3-H4 using anti-H4K12Ac antibodies and cohesin using anti-XRAD21 and anti-XSMC1 antibodies. H3 protein levels are detected using anti-H3 and exhibits a ~90-95% depletion of soluble H3-H4 heterodimers compared to IgG-depleted extract. XSMC1 protein levels are detected using anti-XSMC1, which exhibits a ~90% depletion. We used anti-DM1a that detects tubulin as a loading control. (B) Co-immunodepletion of H3-H4 using anti-H4K12Ac antibodies and condensin using anti-XCAP-C and anti-XCAP-E antibodies. H3 protein levels are detected using anti-H3 and exhibits a ~90% depletion of soluble H3-H4 heterodimers compared to IgG-depleted extract. XCAP-C protein levels are detected using anti-XCAP-C, which displays a ~85% reduction, although the signal is rather weak and made it challenging to quantify. We used anti-DM1a that detects tubulin as a loading control.

## Supplementary video captions

**Movie 1:** Addition of crude *Xenopus* egg extract to a single strand of λ-phage DNA, visualized using Sytox Orange, leads to the generation of multiple highly-enriched DNA clusters, suggestive of nucleosomal formation along the strand.

**Movie 2:** Addition of crude *Xenopus* egg extract to a single strand of λ-phage DNA, visualized using Sytox Orange, leads to the generation of multiple highly-enriched DNA clusters, suggestive of nucleosomal formation along the strand.

**Movie 3:** Example of loop formation in nucleosome-depleted egg extract arrested in metaphase visualized using Sytox Orange. The movie duration is 87 seconds and the scale bar is 5 μm.

**Movie 4:** Example of loop formation in nucleosome-depleted egg extract arrested in metaphase visualized using Sytox Orange. The movie duration is 90 seconds and the scale bar is 2 μm.

**Movie 5:** Example of loop formation in nucleosome-depleted egg extract in metaphase visualized using Sytox Orange. The movie duration is 195 seconds and the scale bar is 2 μm.

**Movie 6:** Example of loop formation in nucleosome-depleted egg extract in interphase visualized using Sytox Orange. The movie duration is 108 seconds and the scale bar is 2 μm.

**Movie 7:** Example of loop formation in nucleosome-depleted egg extract in interphase visualized using Sytox Orange. The movie duration is 80 seconds and the scale bar is 2 μm.

**Movie 8:** Example of loop formation in nucleosome-depleted egg extract in interphase visualized using Sytox Orange. The movie duration is 34 seconds and the scale bar is 2 μm.

**Movie 9:** Example of hydrodynamically stretched loops in nucleosome-depleted extract arrested in metaphase visualized using Sytox Orange. The movie duration is 19 seconds and the scale bar is 5 μm.

**Movie 10:** Example of hydrodynamically stretched loops in nucleosome-depleted extract arrested in metaphase visualized using Sytox Orange. The movie duration is 16 seconds and the scale bar is 5 μm.

**Movie 11:** Example of a hydrodynamically stretched loop in nucleosome-depleted interphase extract visualized using Sytox Orange. The movie duration is 104 seconds and the scale bar is 2 μm.

**Movie 12:** Example of a hydrodynamically stretched loop in nucleosome-depleted interphase extract visualized using Sytox Orange. The movie duration is 4 seconds and the scale bar is 2 μm.

**Movie 13:** Example of loop formation in non-depleted crude extract visualized using Sytox Orange. The movie duration is 87 seconds and the scale bar is 5 μm.

## References

1. Hirano, T. & Mitchison, T. J. Cell cycle control of higher-order chromatin assembly around naked DNA in vitro. J. Cell Biol. 115, 1479–1489 (1991).

2. Rowley, M. J. & Corces, V. G. Organizational principles of 3D genome architecture. Nat. Rev. Genet. 19, 789–800 (2018).

3. Nagano, T. et al. Cell-cycle dynamics of chromosomal organization at single-cell resolution. Nature 547, 61–67 (2017).

4. Bouwman, B. A. M. & de Laat, W. Getting the genome in shape: The formation of loops, domains and compartments. Genome Biol. 16, 1–9 (2015).

5. Bonev, B. & Cavalli, G. Organization and function of the 3D genome. Nat. Rev. Genet. 17, 661–678 (2016).

6. Smith, E. M., Lajoie, B. R., Jain, G. & Dekker, J. Invariant TAD Boundaries Constrain Cell-Type-Specific Looping Interactions between Promoters and Distal Elements around the CFTR Locus. Am. J. Hum. Genet. 98, 185–201 (2016).

7. Dekker, J. & Mirny, L. The 3D Genome as Moderator of Chromosomal Communication. Cell 164, 1110–1121 (2016).

8. Natalia Naumova, Maxim Imakaev, Geoffrey Fudenberg, Ye Zhan, Bryan R. Lajoie, Leonid A. Mirny, J. D. Organization of the Mitotic Chromosome. Science 342, 948–953. (2013).

9. Goloborodko, A., Imakaev, M.V., Marko, J.F. & Mirny, L. Compaction and segregation of sister chromatids via active loop extrusion. Elife (2016).

10. Uhlmann, F. SMC complexes: from DNA to chromosomes. Nat. Rev. Mol. Cell Biol. 17, (2016).

11. Kinoshita, K. & Hirano, T. Dynamic organization of mitotic chromosomes. Curr. Opin. Cell Biol. 46, 46–53 (2017).

12. Sutani, T. et al. Condensin targets and reduces unwound DNA structures associated with transcription in mitotic chromosome condensation. Nat. Commun. 6, (2015).

13. Nasmyth, K. Disseminating the genome: Joining, Resolving, and Separating Sister Chromatids During Mitosis and Meiosis. 673–745 (2001).

14. Alipour, E. & Marko, J. F. Self-organization of domain structures by DNA-loop-extruding enzymes. Nucleic Acids Res. 40, 11202–11212 (2012).

15. Abramo, K., Valton, A., Venev, S. V, Ozadam, H. & Fox, A. N. A chromosome folding intermediate at the condensin-to-cohesin transition during telophase. biorXiv 678474 (2019).

16. Rao, S. S. P. et al. A 3D map of the human genome at kilobase resolution reveals principles of chromatin looping. Cell 159, 1665–1680 (2014).

17. Yatskevich, S., Rhodes, J. & Nasmyth, K. Organization of Chromosomal DNA by SMC Complexes. Annu. Rev. Genet. 53, 1–38 (2019).

18. Fudenberg, G. et al. Formation of Chromosomal Domains by Loop Extrusion. CellReports 15, 2038–2049 (2016).

19. Nuebler, J., Fudenberg, G., Imakaev, M., Abdennur, N. & Mirny, L. A. Chromatin organization by an interplay of loop extrusion and compartmental segregation. Proc. Natl. Acad. Sci. U. S. A. 115, E6697–E6706 (2018).

20. Ganji, M., Kim, S. H., Van Der Torre, J., Abbondanzieri, E. & Dekker, C. Intercalation-based single-molecule fluorescence assay to study DNA supercoil dynamics. Nano Lett. 16, 4699–4707 (2016).

21. Yan, J. et al. Micromanipulation Studies of Chromatin Fibers in Xenopus Egg Extracts Reveal ATP-dependent Chromatin Assembly Dynamics. MBoC 18, 464–474 (2007).

22. Gruszka, D. T., Xie, S., Kimura, H. & Yardimci, H. Single-molecule imaging reveals control of parental histone recycling by free histones during DNA replication. bioRxiv 789578 (2019).

23. Zierhut, C., Jenness, C., Kimura, H. & Funabiki, H. Nucleosomal regulation of chromatin composition and nuclear assembly revealed by histone depletion. Nat. Struct. Mol. Biol. 21, 617 (2014).

24. Ganji, M. et al. Real-time imaging of DNA loop extrusion by condensin. Science 360, 102–105 (2018).

25. Banigan, E. J. & Mirny, L. A. Limits of chromosome compaction by loop-extruding motors. PRX 9 031007 (2019).

26. Banigan, E. J., Berg, A. A. Van Den, Brandão, H. B. & Marko, J. F. Chromosome organization by one-sided and two-sided loop extrusion. bioRxiv 815340 (2019).

27. Terence R. Strick, Tatsuhiko Kawaguchi, T. H. Real-time detection of single-molecule DNA compaction by condensin I. Curr. Biol. 128, 189–190 (2004).

28. Muwen Kong, Erin Cutts, Dongqing Pan, Fabienne Beuron, Thangavelu Kaliyappan, Chaoyou Xue, Ed Morris, Andrea Musacchio, Alessandro Vannini, E. C. & Greene. Human condensin I and II drive extensive ATP – dependent compaction of nucleosome – bound DNA. biorXiv 683540 (2019).

29. Losada, A., Hirano, M. & Hirano, T. Identification of Xenopus SMC protein complexes required for sister chromatid cohesion. Genes Dev. 12, 1986–1997 (1998).

30. Rao, S. S. P. et al. Cohesin Loss Eliminates All Loop Domains. Cell 171, 305–320 (2017).

31. Schwarzer, W. et al. Two independent modes of chromatin organization revealed by cohesin removal. Nature 551, 51–56 (2017).

32. Davidson, I. F. et al. Rapid movement and transcriptional re-localization of human cohesin on DNA. 35, 2671–2685 (2017).

33. Abdennur, N. et al. Condensin II inactivation in interphase does not affect chromatin folding or gene expression. bioRxiv 437459 (2018).

34. Hirano, T., Kobayashi, R. & Hirano, M. Condensins, Chromosome Condensation Protein Complexes Containing XCAP-C, XCAP-E and a Xenopus Homolog of the Drosophila Barren Protein. Cell 89, 511–521 (1997).

35. Sanborn, A. L. et al. Chromatin extrusion explains key features of loop and domain formation in wild-type and engineered genomes. Proc. Natl. Acad. Sci. 112, E6456–E6465 (2015).

36. Ren, G. et al. CTCF-Mediated Enhancer-Promoter Interaction Is a Critical Regulator of Cell-to-Cell Variation of Gene Expression. Mol. Cell67, 1049–1058 (2017).

37. Hansen, A. S., Pustova, I., Cattoglio, C., Tjian, R. & Darzacq, X. CTCF and cohesin regulate chromatin loop stability with distinct dynamics. Elife 6, 1–33 (2017).

38. Murray, A. W. Cell cycle extracts. in Methods in cell biology 36, 581–605 (Elsevier, 1991).

39. Jenness, C., Wynne, D. J. & Funabiki, H. Protein immunodepletion and complementation in Xenopus laevis egg extracts. Cold Spring Harb. Protoc. (2018).

40. Groen, A. C., Nguyen, P. A., Field, C. M., Ishihara, K. & Mitchison, T. J. Glycogen-supplemented mitotic cytosol for analyzing Xenopus egg microtubule organization. in Methods in enzymology 540, 417–433 (2014).

41. Smith, S. B., Cui, Y. & Bustamante, C. Overstretching B-DNA: the elastic response of individual double-stranded and single-stranded DNA molecules. Science 271, 795–799 (1996).

42. Marko, J. F., Siggia, E. D. & Smith, S. Entropic Elasticity of X-Phage DNA Explicit and Implicit Learning and Maps of Cortical Motor Output. Science 265, 1599–1600 (1994).

